# The effect of surface properties on the interactions of particles and marine mucous filters

**DOI:** 10.1101/2025.01.21.634085

**Authors:** Yuval Jacobi, José A. Epstein, Ariella Paz, Uri Shavit, Gitai Yahel, Guy Z. Ramon

**Affiliations:** Faculty of Civil and Environmental Engineering, Technion – Israel Institute of Technology; Faculty of Marine Sciences, Ruppin Academic Center

## Abstract

Free-living suspended cells form the foundation of marine food webs, making suspension feeding a key mode in aquatic ecosystems. Suspension feeders across phyla use diverse filtration mechanisms, often relying on mucus and low-pressure pumps with high filtration efficiencies and self-cleaning capabilities. Traditionally, particle capture was thought to depend on size alone, but recent evidence highlights the importance of physicochemical surface interactions between prey cells and filtration apparatuses. In this study, we investigated the capture of 0.3-3 µm particles by ascidians and found that coating particles with amphiphilic polymers altered their mobility within the mucous filter, affecting capture efficiency. Surface interactions, such as steric repulsion, significantly influence particle mobility and inversely correlate with capture success. Furthermore, we discovered that the mucous filter in ascidians is much thicker (∼5 µm) than previously believed, functioning as a continuous sheet. These findings suggest a need to reevaluate suspension feeding models, with implications for marine ecosystems and filtration technology development.

## Introduction

Feeding on suspended marine microorganisms is a common trophic strategy, named “suspension feeding”. The capture of particles by suspension feeders is often described as a process made of three consecutive steps: the *encounter* of prey particles with the capture apparatus, the *retention* of prey particles, and the *handling* of retained particles (1–3). At the microscale, the capture systems are described as arrays of cylinders that collect food particles from low-velocity (i.e., with Reynolds number lower than one) flows (3–6).

The involvement of mucus in suspension-feeding is widespread (7). Among the most prominent methods is *mucus-net filter-feeding,* which is found in tunicates such as salps, appendicularians, and ascidians. These suspension feeders drive their feeding currents through a mucous filter which retains the food particles and transports them toward the digestive tract (8, 9). *Mucus-net filter-feeding* is also used by some gastropods, polychaete worms, and lancelets (10). The gills of bivalves, which act as a particle capture system, contain mucus that transports and possibly retains particles (11–13). It is thought that some sponges have a mucus-like layer around the collar filter that is involved in particle retention (14, 15). Corals, as well as jellyfish, utilize mucus for many purposes, among which is particle capture (10). Adhering to the “textbook” description of *mucus-net filter feeding* suggests that mucus is involved in both *encounter* and *retention* during particle capture by these animals. For other suspension feeders (e.g., bivalves, pteropods, corals, and jellyfish) mucus is most likely involved in particle *retention* by its adhesive properties and particle *handling* via mucociliary transport.

Due to their relatively simple structure, ascidians (sea squirts) are excellent models for studying particle capture mechanisms with mucus. The feeding current of ascidians is unidirectional: from the inlet, through a mucous filter, and out to the sea. Adult solitary ascidians are usually a few centimeters long, and their body has two openings: the inhalant and exhalant siphons. Ascidians draw water through the inhalant siphon into a perforated organ known as the branchial sac, which is where filtration occurs. The inner walls of the branchial sac pores are covered with ciliated cells that function as a shear pump and force water in and through the mucous filter. On the downstream side of the branchial sac, there is a cavity known as the atrium through which the filtered water passes and then exits to the sea via the exhalant siphon (16). The gland that secretes the mucous filter is called the endostyle. Opposite to the endostyle, there is a structure known as the dorsal lamina. As illustrated in Fig. 1, when ascidians feed, the endostyle constantly produces mucus that is draped on the inward-facing surface of the branchial sac. This mucous filter is constantly propagated by an additional set of cilia that serve as a conveyor belt that moves the filter along the inner circumference of the brachial sac towards the dorsal lamina (8, 17). At the dorsal lamina, the mucous filter, along with the retained particles it collects, is drawn into the digestive tract (9).

**Figure 1.**
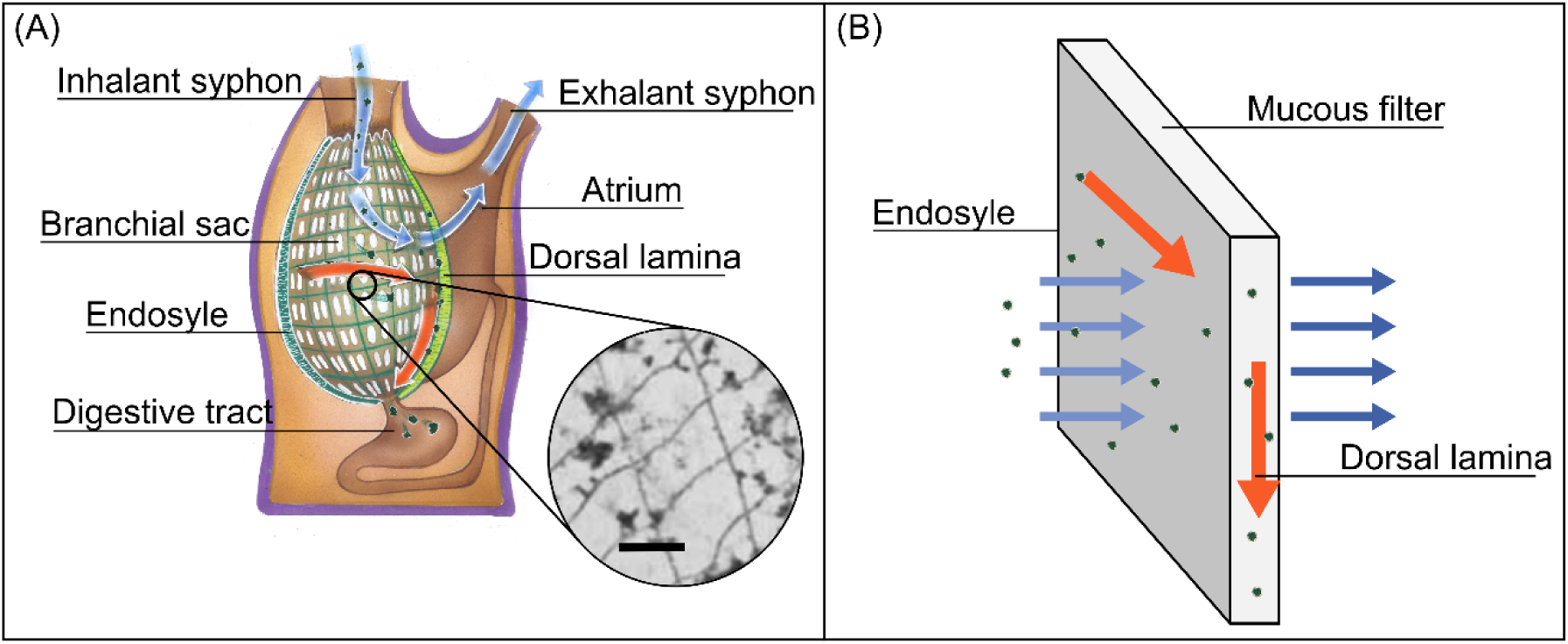
(A) - Drawing of a longitudinal section of a solitary ascidian. Water (blue arrows) enters the branchial sac through the inhalant siphon along with food particles (dark green shapes). The water then leaves through the openings of the branchial sac (white ellipses) and out to sea. Food particles are retained by the mucous filter that lines the inner side of the branchial sac. The mucous filter is constantly secreted from the endostyle and conveyed towards the dorsal lamina (red arrows represent mucus movement). At the dorsal lamina, mucus and food particles are drawn into the digestive tract. The mucous filter is thought to be a thin rectangular mesh, as can be seen in the inset of a transmission electron microscopy image of the mucous filter. This image is of the mucus of the ascidian Ciona intestinalis. The scale bar in the inset is 1 µm. Adapted with permission from (18). (B) – A schematic illustration of the mucous filter. Water carries food particles towards the mucous filter, which is continuously secreted by the endostyle and collected at the dorsal lamina. The mucous filter is continuously secreted and moved toward the dorsal lamina (red arrows) acting as a conveyor belt that transports the particles that are embedded in it to the dorsal lamina and eventually the digestive tract.

The “textbook” description of the mucous filter in ascidians is a mesh made of cylindrical mucous fibers that are 10–40 nm thick (Fig. 1A). These fibers are thought to be organized into a rectangular mesh with pores that are 0.2–0.5 μm wide and 0.5–2.2 μm long and with little variation between species (8, 16, 18). Ascidians were shown to efficiently capture particles as small as 0.3 µm (19, 20), but surprisingly, microbial plankton is captured by ascidians less efficiently than polystyrene particles of similar sizes (21). Furthermore, studies show that some bacterial clades can evade capture by ascidians due to their surface properties rather than their size or shape (19, 22). There are no known mechanisms of selection or sorting during particle capture in ascidians. Surprisingly, it was also found that planktonic microbes that are large enough to be retained by simple sieving (i.e., cells that are larger than the pore size) were captured at less than 100% efficiency (∼60-70%, (21)). These observations cannot be easily reconciled with the notion that the ascidian filter is a thin rectangular mesh-like structure made of nano-scale mucous fibers that capture food particles by simple sieving.

In the present work, ascidians were used as a model to study particle capture (and evasion) by mucous filters. We studied particles interacting and transporting through the mucus and the effect of surface interactions on particle capture using several complementary approaches. First, the surfaces of artificial micro-particles were modified and characterized. This modification was done by adding a polymer brush to the surface (23) of polystyrene microspheres (hereafter beads). Particles coated with polymer brushes are less prone to aggregate or adsorb to surfaces (24). The beads were coated with long-chain tri-block copolymer surfactants commercially known as Poloxamers or Pluronics. Pluronics are made of a poly(propylene oxide) (PPG) core, which absorbs on the bead’s surface, and two poly(ethylene glycol) (PEG) chains extending away from the interface (25). Four different Pluronic types were used, all with relatively similar PPG contents but with significantly different lengths of their PEG chains. This resulted in five types of experimental beads, referred to hereafter as P0, P0.3, P1.2, P6.8, and P12, respectively. The purpose of this nomenclature is to establish a reference to the PEG content of the coated beads and, hence, the relative thickness of the polymer brush layer. Interaction and transport of the beads through the mucus were studied by 1. In-vivo measurement of particle capture efficiency; 2. Measurement of the diffusive and advective transport of beads through the mucous filter; and 3. Characterization of the mucous filter by cryogenic scanning electron microscopy (cryo-SEM) and proteomic analysis. Results showed that beads coated with polymer brushes of intermediate thickness are captured less efficiently than other beads. Furthermore, our findings contradict the predictions of the accepted model of the ascidian filter as a two-dimensional array of thin mucous cylinders. Based on these observations, we suggest an alternative model: a thick, continuous mucous sheet through which the unretained particles are transported at different rates according to their surface properties. Our results highlight the important role that surface properties play in the way colloids interact with mucus and suggest a potential mechanism for some of these interactions.

## Results and discussion

### The effect of bead size and surface properties on particle capture

Regardless of their surface modifications, the capture of large beads (diameter of 3 and 10 µm) by the mucous filter of all ascidian species we tested approached 100% efficiency (efficiency defined as the ratio of the number of captured particles to the number of all particles of the same size and type inhaled by the animal). Smaller coated beads (diameters less than 1 µm) were captured at a significantly lower efficiency than the uncoated (P0) particles (Fig. 2 A-D). Altering the length of the PEG chains on the surface of the beads had a significant but non-monotonic effect on the capture efficiency by the ascidians *H. momus* and *M. exasperatus* (Fig. 2 E-F). Beads coated by Pluronic molecules with intermediate length of the PEG chains were captured less efficiently than uncoated particles or particles coated by short (0.3 KD) and long (12 KD) PEG chains (ANOVA with permutations, *p* < 0.001, n=15, Fig. 5, Table S2). It is speculated that the low capture efficiency of coated beads, especially those with intermediate PEG content, is a consequence of steric repulsion between the mucus filter and the particle surface induced by the presence of the polymer-brush coating. This assertion is strengthened by the fact that particles coated with a PEG brush were previously shown to penetrate human reproductive mucous layers better than uncoated particles, and their penetration was similarly altered by the length of the PEG brush, resembling the effect of polymer brush length on the capture efficiency by ascidians (26).

**Figure 2.**
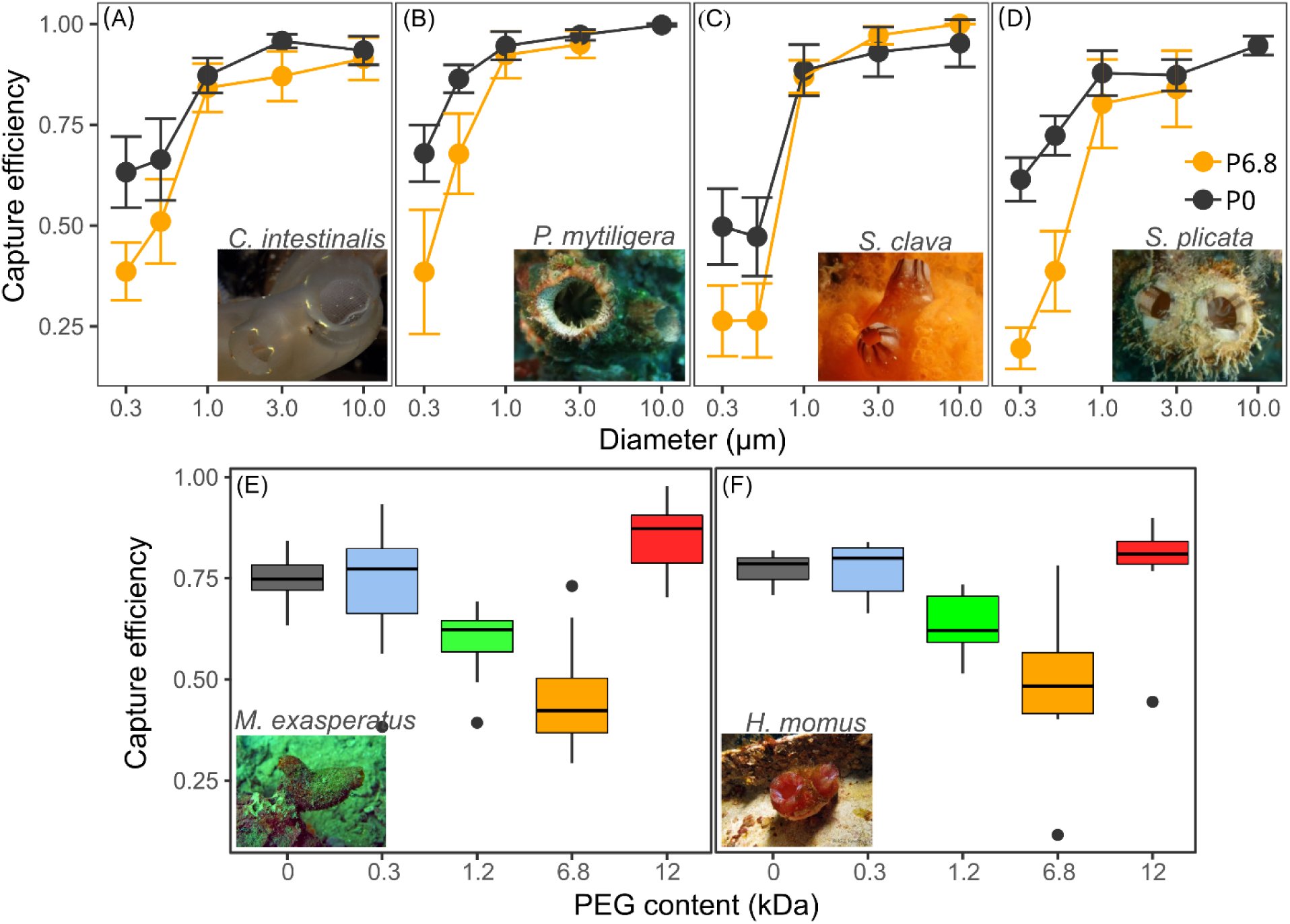
(A-D) - Capture efficiency of coated beads (P6.8, orange) and uncoated beads (P0, black) by the ascidians C. intestinalis (A), P. mytiligera (B), S. clava (C), and S. plicata (D). Error bars are 95% Confidence interval of the mean (CI). Sample size: (A) P0 n=17, P6.8 n=17; (B) P6.8: n=9, P0: n=26; (C) P0 n=9, P6.8 n=9; and (D) P6.8: n=13, P0: n=20. (E, F) - Capture efficiency of 0.5 µm P0, P0.3, P1.2, P6.8, and P12 beads (gray, blue, brown, orange, and red boxes, respectively) by the ascidians M. exasperatus (E) and H. Momus (F). The median is presented as a bolded horizontal line, the box encompasses the second and third quartiles, and the whiskers extend to the nearest point that is < 1.5 times the interquartile range. Circles are outliers. H. momus n=9, M. exasperatus, n=10. Photos of these species are shown in the bottom right of each plot. Photos by M. Gewing (A), N. Shankar (B), and G. Koplovitz (D, F), and courtesy of http://ascidians.com/ (C, E).

### Surface properties influence transport characteristics through mucus

To study the diffusive transport of the beads, we measured their mean square displacement (MSD) and used the slope of the time-scale-dependent MSD curve (*MSD*(*τ*)) as an estimate of their diffusivity through different media. When suspended in water, the measured *MSD*(*τ*) curves of all beads overlapped, indicating that the beads’ diffusivity is nearly identical regardless of their coating. This means that the different coatings had a negligible effect on the hydrodynamic size of the beads (Fig. 3A). In contrast when suspended in mucus, the diffusive mobility of beads with different coatings was significantly impacted by surface modifications (Fig. 3B). The *MSD*(*τ*) curves of the different bead types indicate that, except for P12, mobility within the mucus increased with chain length (Fig. 3 A-B).

**Figure 3.**
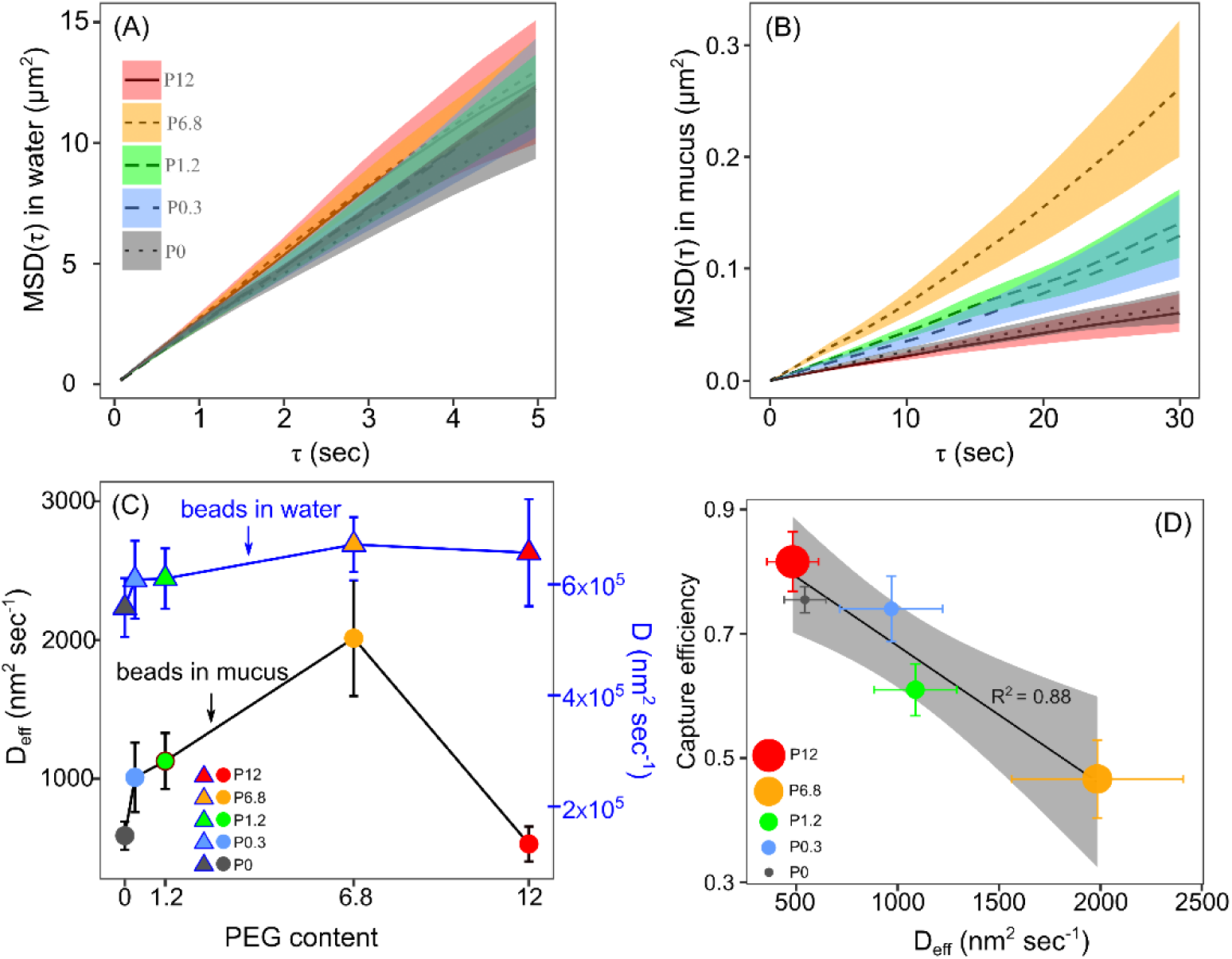
(A, B) – The *MSD*(*τ*) curves of the experimental beads in distilled water (A) and in fresh mucus harvested from H. Momus (B). Shaded areas show the 95% confidence interval for the MSD in each lag time, τ. (C) - The diffusion coefficients as a function of the PEG content in mucus (circles and black line, left axis) and in distilled water (triangles and blue line, right axis). Sample size of the MSD measurements is 38, 44, 47, 49, and 55 in mucus, and 53, 61, 51, 104, and 25 in water for the P0, P0.3, P1.2, P6.8, and P12 beads, respectively. Error bars represent the 95% CI. (D) - The capture efficiency of the experimental beads by H. momus (n=9) as a function of Deff. The diameter of the circular markers is proportional to 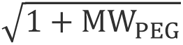 where MW_PEG_is the PEG molecular weight. The black line is the linear fit for these data and the shaded area shows the 95% CI for the regression, with annotation of the coefficient of determination (R2). Error bars are 95% CI.

Within the time frame of the performed experiments (30 seconds), the *MSD*(*τ*) curves of all beads in mucus exhibited a monotonic increase (Fig. 3B). The effective diffusion coefficient (D_eff_) for each bead type could thus be estimated by taking the derivative of *MSD*(*τ*) according to Eq. (S2a) (see supplementary material). Beads coated with Pluronics containing intermediate PEG chains had a relatively high D_eff_ (1127 ± 202 and 2013 ± 418 nm^2^ sec^-1^ for P1.2 and P6.8, respectively), compared to beads coated by Pluronics with smaller chains (P0.3, 1010 nm^2^ sec^-1^, Fig. 3C). Beads coated with the largest PEG chains (P12) had a D_eff_ comparable to that of the uncoated beads (529 ± 126 and 589 ± 102 nm^2^ sec^-1^ for P0 and P12 respectively). Notably, the effective diffusion coefficient accounted for >98% of the variance in capture efficiency by the tested ascidians, and the two parameters were negatively correlated (Spearman r = -0.96, p = 0.016), meaning that beads with a higher effective diffusion coefficient are less efficiently captured (Fig. 3D).

Filtration experiments with fresh mucus harvested from live ascidians revealed that, in concurrence with the MSD results, P6.8 beads (which also showed the highest D_eff_) travel through the mucus faster than uncoated (P0) and P12 beads (Fig. 4A, 4B, and S3). P6.8 beads accumulated at the mucus-supporting membrane interface (at z = 0, as illustrated in Fig. 4E) faster than uncoated beads (Fig. 4C). Plotting the distance at which coverage reached half of the maximum value (hereafter referred to as Z_50._ The maximum concentration was measured at z = 0) vs. time, enabled an estimate of the average velocity of the beads across the mucous layer, under the experimental conditions. The velocities measured for uncoated and P6.8 beads (0.2 and 0.6 µm min^-1^, respectively) were at least three orders of magnitude lower than the estimates found in the literature of the average velocity of water crossing through the mucous filter (∼0.4 mm min^-1^, 127, 28).

**Figure 4.**
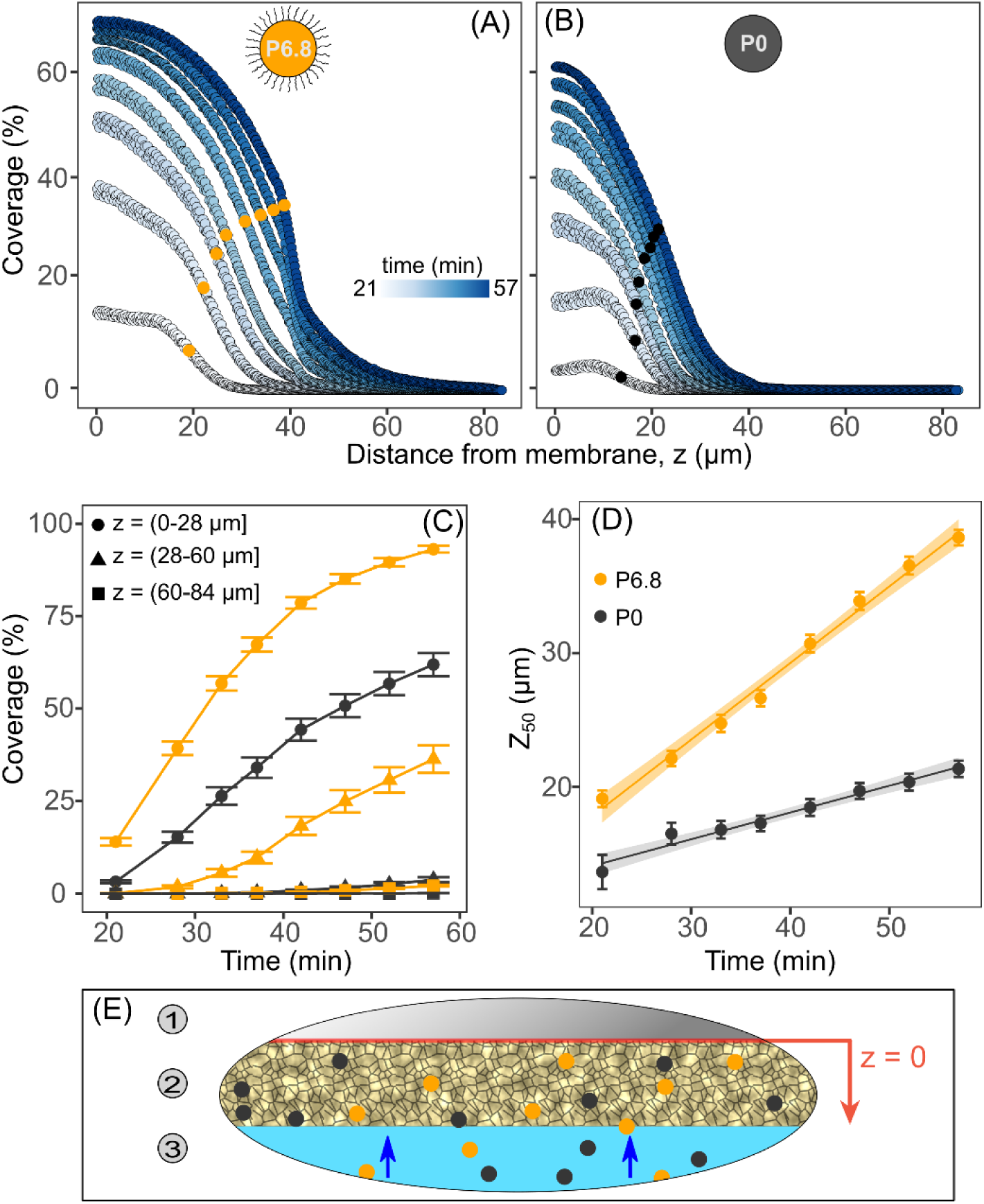
(A, B) - The relative coverage of 0.5 µm P6.8 (A) and P0 beads (B) in the mucus at different distances from the supporting membrane. The time that has passed since beads were introduced to the system is marked by different colors. The points where coverage is equal to half of its maximal value on each curve are marked by orange (A) and gray circles (B). (C) - The mean relative coverage of particles at different distances from the supporting membrane as a function of the time. Coverage values are presented as the average over the range of distances from the supporting membrane at 4 adjacent visualized areas. The distance range is presented by different orange (P6.8) and gray (P0) markers: Circles 0-28, triangles 28-60, and squares 60-84 µm. Error bars are 95% CI. (D) – Z_50_ (the z value of the orange and gray circles marked on A and B) as a function of time. P0 particles (gray circles) are slower than P6.8 particles (orange circles). Lines are the linear fit for these data, and the shaded area shows the 95% CI for the regression. For the P6.8 z(µm) = 0.6 t(minute) + 6 and R^2^ = 0.99; for the P0 z(µm) = 0.2 t(minute) + 10 and R^2^ = 0.97. Error bars are the 95% CI, n=4. (E) - Schematic illustration of the experimental system. 1. supporting membrane, 2. mucus specimen, and 3. the feed solution. Particles (orange and gray circles) were introduced to the flowing water at t = 0 and are carried by the permeate flux (blue arrows) through the mucus until they deposit at the mucus-membrane interface at z=0 (marked with a red line).

The correlation between particle transport through the mucus and their capture efficiency shows that the surface properties of particles and, presumably, the physicochemical surface interactions with the mucous filter affect the mobility of particles through the filter and, most likely, influence capture. The main conclusion is intuitive: more mobile particles cross the mucus faster than the less mobile particles.

### Proposed mechanism: polymer brushes, steric repulsion, and entanglements

Polymer brushes, formed by the PEG on the surface of the particles, induced a repulsive steric force that countered attractive interactions (e.g., hydrophobic interactions and London-van der Waals forces) between the coated beads and the mucus in the capture apparatuses of ascidians. The polymer brush on the beads’ surface is made of adsorbed PEG, and as the PEG content increased (from 0, through 0.3, 1.2, to 6.8 kDa), capture efficiency decreased. However, the capture efficiency of particles coated with longer PEG chains (12 kDa) is as high as that of uncoated beads. In a brush, the average distance between neighboring chains tethered to the beads increases with distance from the surface of the beads due to particle curvature (25). For a thick enough polymer layer, the conformation at the free end of the tethered chains may not be the stretched conformation associated with polymer brushes, and some parts of the chain may assume a relaxed conformation (23). This kind of structure, where the bases of the tethered polymers are stretched, and the free ends are not, is often referred to as a polymer mushroom (29). The high capture efficiency of P12 beads and their lower MSD (Figs. 3 and 4, and Table S2) may be a result of entanglements of the relatively long polymers with the mucus, made possible by their supposed mushroom structure.

In marine suspension feeding, electrostatic interaction between particles and filters is usually neglected because of the high ionic strength of seawater (5). The results of this study agree with the assertion that surface charge has little influence on capture. The measurements of the ζ potential of the experimental beads showed that P0 beads carry a substantial (negative) surface potential (∼ -25 mV), while the other beads are characterized by uniformly low ζ potential. The ascidian mucus is believed to be negatively charged as well (Flood & Fiala-Medioni 1981), but P0 particles were captured with higher efficiencies than particles that carry less surface charge (e.g., P6.8 in Fig. 3).

Polymer-brush-induced steric repulsion is largely independent of the chemical makeup of the polymers in the brush (23). Steric repulsion will occur if the fluid in which the particles are suspended is a good solvent for the polymers and if the grafting density of the polymer chains on the particle’s surface is high enough (i.e., the distance between grafting points is smaller than the radius of gyration of the polymer in its ideal chain conformation (23)). These findings, therefore, suggest a potential mechanism for the many observations (19, 30, 31) of the effect of surface properties on particle capture. Furthermore, interactions of microbes and other particles with mucus are a widespread phenomenon in the ocean, occurring in processes such as the formation of phycospheres, transparent exopolymer particles, and other types of marine organic aggregates. These aggregates are crucial in transporting matter and energy to the deep sea. Steric repulsion due to surface polymer brushes may also be involved in these aggregation processes and play an essential role in such fluxes.

### Revisiting the structure of the ascidian filter

Early studies described the mucous filter as a thick mucus sheet (32). Many years later, Flood and Fiala-Medioni (18) observed fixed, dried, and gold-palladium-coated samples of the mucous filter of several ascidian species with transmission and scanning electron microscopy. They reported that the ascidian mucous filter is a submicron-sized rectangular grid made of thin (∼40 nm) mucous fibers (Fig. 1A), and their observations were later corroborated by others that used similar techniques (33, 34). Stemming from these observations is the notion, adopted by all invertebrate textbooks, that the structure of the mucous filter of ascidians is an almost two-dimensional, submicron rectangular mesh (9, 16, 35). However, as discussed in the following sections below, this description is hard to reconcile with current observations.

Rather than a two-dimensional thin mesh, the findings of this work suggest that the mucous filter of ascidians may be described as a thick layer of mucus continuously secreted by the endostyle and propagated at a velocity of ∼1 cm m^-1^ (17 and references therein) towards the dorsal lamina where the filter and embedded particles are drawn to the digestive tract. A volumetric water flow enters the filter with the particles it transports; thus, all particles transported by the feeding current *encounter* the mucus. This water flow is responsible for the particle advection across the mucus layer. At the downstream side of the filter, the water leaves the filter out to sea with the particles that were not captured. Particles trapped by the mucus are conveyed with the mucus, arriving at the dorsal lamina within a few minutes (17). Particles that reach the dorsal lamina before reaching the downstream side of the filter are captured, while particles that reach downstream before they are conveyed to the dorsal lamina escape (Fig 1B). If the mucus filter is a thick and continuous hydrogel-like layer, particles that traverse the thick mucus fast enough can evade capture by crossing to the downstream side of the filter before the mucus carries them to the dorsal lamina and the digestive tract (see Fig. 1). Thus, the capture efficiency of ‘fast’ particles with high motility is lower than that of ‘slow’ ones. It should be noted that in the diffusive and advective transport experiments, the mucus was likely not in the same configuration as it is on the branchial basket.

The surface properties of the beads affect the rate of their diffusion (Fig. 3) and advection (Fig. 4) through the mucus. These observations do not necessarily contradict the possibility that the filter is made of a thin mesh, and the surface modifications may influence the bead transport and the retaining force needed for capture. However, unlike the expected capture of a true ‘mesh’, there is no strict size-based cutoff – large particles can avoid capture if they traverse the filter fast enough.

### Mucus extraction yielded more mucus than expected

The mucus harvested from *H. momus* individuals (Fig. 5A. 49 ± 25 µL per ascidian, mean ± 95% CI, n=10 pooled mucus samples from 113 ascidians) was in much higher quantities than expected based on published mesh thickness. These values are supported by the reported mean (± 95% CI) dry weight of a single mucous mesh of 2.4 ± 0.6 mg (17). The ascidians we used for mucus harvesting were relatively large individuals having a branchial sac with an average area of 10.3 ± 1.2 cm^2^ (17). If the entire area of the branchial sac is covered by a mucous mesh and its thickness is 50 nm, as previously suggested (18, 33, 34), the volume of the entire mucous filter should be no larger than 0.05 µL, three orders of magnitude smaller than the observed values. Based on this simple measurement and the above considerations, it is concluded that the ascidian mucus filter is ∼3 orders of magnitude thicker than the proposed ∼50 nm.

**Figure 5.**
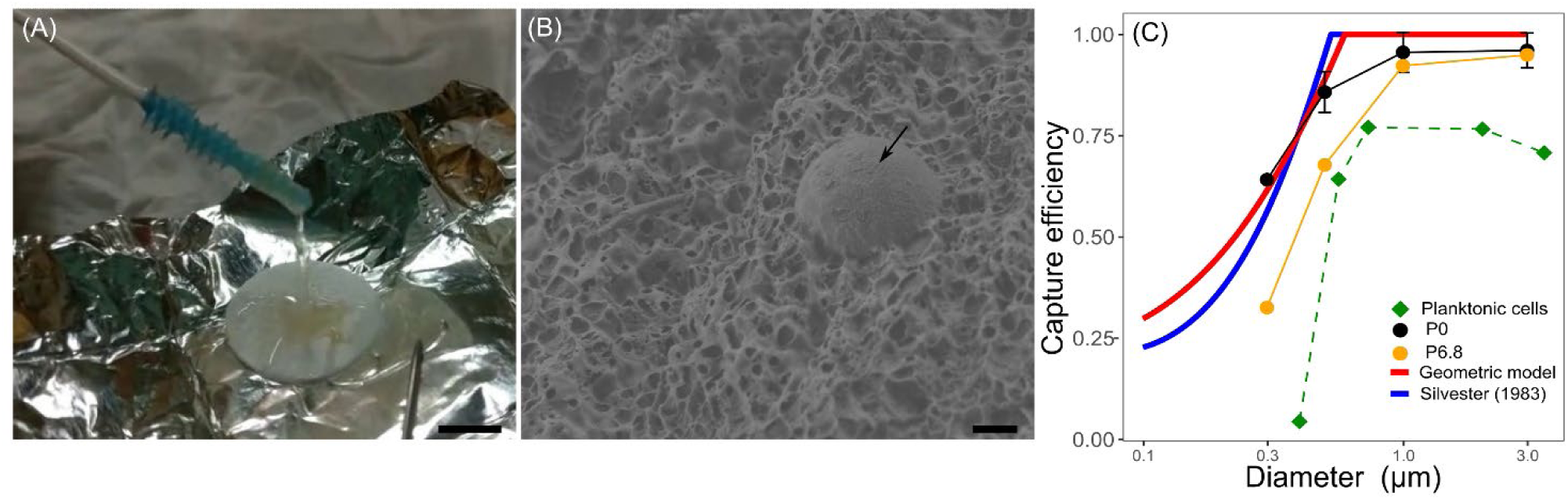
(A) - A mucus sample harvested with a swab from the ascidian *H. momus*. The scale bar is 10 mm. (B) - Cryo-SEM image of a 3 μm P0 bead (black arrow) embedded in the mucus. The structure of the mucus appears to be like that of a heterogeneous hydrogel. The scale bar is 1 μm. For specimen preparation, see the supplementary material (surface imaging methodology). (C) - Observed and calculated capture efficiencies of cells and beads by the studied ascidians. The model suggested by Silvester (1983) (Blue line) and the Geometric model developed in this work (red line, see supplementary material) were fitted to the observed capture efficiency for uncoated beads (black circles) by adjusting the pore size of the modeled mesh. Error bars are omitted for clarity (but are presented in Fig. S8). For planktonic cells n=5, for P0 n=5, and for P6.8 n=4 ascidian species.

### Microscopy of the mucous filter

Observations of fresh specimens of the brachial sac of *H. momus* and *S. plicata* with mucus were done using cryo-SEM (Fig 5B and Fig S4). These images corroborated the hypothesis that the mucous filter is thicker than previously thought, and its structure resembles that of a heterogeneous hydrogel, not a highly organized thin mesh. Unlike the electron microscopy methods used to observe the mucous filter in the past (18, 33, 34), preparing specimens for cryo-SEM does not require manipulations other than rapid freezing in liquid ethane. It should be noted that the specimens observed by cryo-SEM were hydrated with seawater; thus, brine facets, typical artifacts of the method, were visible in most samples. Moreover, specimens small enough for an optimal freezing rate were hard to obtain, and the use of larger specimens resulted in the formation of ice crystals that disrupted large regions of specimens. Therefore, observations of the uninterrupted structure of the mucus could be done only on the specimen’s edges. Due to these technical difficulties, it is hard to draw quantitative conclusions from the cryo-SEM images. Nevertheless, these images do suggest that the mucous filter resembles a heterogeneous hydrogel and provide no evidence for the existence of a thin, organized mesh.

### Proteomic analysis

The amount of protein found in mucus samples was minuscule (data not shown), and the total N content of the mesh was 4.4±0.7%. No peptides unique to the mucus were found in the proteomics analysis, and the peptide composition of the tissue and mucus samples was not significantly different (ANOSIM, *p*≥0.1), regardless of transformation or similarity indices used. These observations suggest that the mucus is not made of some “special” proteins capable of self-assembly but rather a matrix of tangled polysaccharides or other polymers with or without cross-linking bonds.

### Observed capture efficiencies deviate from theoretical predictions of capture by a thin mesh

If the mucous filter is a thin mesh, large particles are retained by mechanical sieving and are expected to be captured at 100% efficiency regardless of their surface properties (36). However, previous studies showed that the capture efficiency of planktonic cells by ascidians is lower than that of similarly sized polystyrene particles not only for particles smaller than the suggested pore size of the mucous mesh (∼0.5 µm) but, surprisingly, also for particle size much larger than the putative mesh pores (3-10 µm, (21)). As explained above, for particles smaller than the pores of the proposed thin mesh model, surface interactions with the mesh may influence particle retention efficiency, e.g., by affecting the retaining adhesive force that keeps particles from rolling off the fibers. However, for particles that are larger than the pore size, this explanation does not hold. An exception could be a case where the large nanoplankton that evade capture are thin and elongated cells with an aspect ratio in the order of 10 that can slip through the rectangular submicron pores (22). However, measurements of the plankton in the water filtered by the ascidians indicate that such cells are extremely rare (21).

As depicted in Fig 5C, theoretical predictions of particle capture with a rectangular mesh (37) are at odds with data on the capture of planktonic cells by ascidians. The discrepancy between the lower-than-expected 100% capture efficiency of large microalgae and beads relative to the proposed dimensions of the ascidian filter was highlighted in previous studies (21) by calculating the probability of particle encounter with fibers of the mesh using the model suggested by Silvester (1983) that is frequently used to predict particle capture by rectangular meshes (20, 27, 38–40). One of the assumptions in the Silvester model is that the particles are considerably smaller than the mesh fiber diameter. However, the published diameter of the ascidian filter fibers is ∼10-50 nm (18, 33, 34), much smaller than the smallest planktonic cells (all larger than 0.2 µm). To account for the potential weakness of the Silvester model, the expected capture efficiency of particles that are smaller than the pore size was also calculated here using a “Geometric model”. This model is based on the proposed geometry of the ascidian rectangular mesh and the size exclusion of passing particles (Fig 5). The residual sum of squares of the observed data from the fitted model was 0.005 for the geometric model and 0.018 for the Silvester (1983) model. However, the geometric model also fails to predict the capture efficiency data of planktonic cells and P6.8 beads by ascidians (Fig 5C).

## Concluding remarks

The surface properties of particles influence the efficiency by which they are captured by the mucous filter, suggesting that the evasiveness of marine phytoplankton and bacteria (see also (19, 21, 30) and references therein) is a consequence of surface interactions. It seems that surface interactions play an important role for suspension feeders that use mucus for particle capture. Specifically, steric repulsion, induced by a coat of polymer brushes on the surfaces of particles, reduces particle capture by mucous filters. Steric repulsion is, to a large extent, independent of the exact chemical makeup of the polymers in the brush layer (23). Polymer brushes and steric repulsion influence bacterial and algal adhesion (41, 42) suggesting that steric repulsion may be an important mechanism in grazing resistance and the formation of marine aggregates. To test this idea, future work should focus on examining the effect of other types of brush layers and on observations of naturally occurring brush-covered and uncovered microbes. This work suggests that in addition to the known mechanisms of grazing resistance exhibited by marine microbes, such as toxicity, chemical deterrents, and extra-cellular shells like the frustules of diatoms and the theca of dinoflagellates, there is likely another mechanism, namely, surface interactions that enable some microbial plankton to escape their predators.

The results presented here call for a revision of our understanding of the structure of the ascidian filter and emphasize the importance of surface interactions, and specifically steric repulsion, in determining the fate of particles that encounter mucus. Furthermore, the methodological protocols established here can help advance studies on the role and mode of action of surface interactions in marine suspension feeding and the formation of organic aggregates.

## Materials and methods

### Experimental particles

#### Surface modification

Polystyrene beads (Fluoresbrite® YG Carboxylate Microspheres, Polysciences) were modified by adding a polymer brush on their surface (23). This was achieved by coating the beads with long-chain tri-block copolymer surfactants commercially known as Poloxamers or Pluronics that are made of poly(ethylene glycol) (PEG) and poly(propylene oxide) (PPG). Four different types were used (Sigma-Aldrich (USA)): L81 (cat. 435430), L64 (cat. 435449), F68 (cat. 412325), and F108 (cat. 542342). The PPG content of Pluronics ranges from 1.7 to 2.6 kDa, while their PEG content varies significantly from 0.3, 1.2, 6.8, and 12 kDa for L81, L64, F68, and F108, respectively. This resulted in five types of experimental beads: uncoated, L81, L64, 68, and F108 coated polystyrene beads, referred to hereafter as P0, P0.3, P1.2, P6.8, and P12, respectively. The purpose of this nomenclature is to establish a reference on the PEG content of the coated beads and, hence, the relative thickness of the polymer brush layer.

#### Surface coverage

To coat the beads, each of the Pluronics was dissolved in water and the mixture was incubated at room temperature on an orbital shaker overnight. A colorimetric method (43) was used to measure the Pluronic concentrations based on the formation of a water-insoluble complex of the Pluronic with cobalt (II) thiocyanate. The Pluronic-cobalt complex is then dissolved in acetone, and the absorbance of the solution at 624 nm is proportional to the Pluronic-cobalt concentration. To measure the amount of adsorbed Pluronic on the surface of the beads, the Pluronic solution was sampled before and after the addition of beads (and an overnight incubation). The second sample was filtered using a 0.2 µm syringe filter and the filtrate was collected. The concentrations of Pluronic in the first and second samples were measured using the colorimetric method, and the concentration of adsorbed Pluronic was calculated as the difference between the two. Results indicated that the average grafting density of the Pluronic molecules on the beads’ surface was between 0.65 and 4 molecules nm^-2^. Additional details of the colorimetric assessment of the grafting density may be found in the supplementary material.

#### Surface hydrophobicity and zeta potential

The hydrophobicity of the beads was evaluated using the “captive bubble” method and a drop shape analyzer (see supplementary material). The presence of the Pluronic coating was verified by the decrease in the magnitude of the ζ potential (measured in 12.3 ppm water and pH = 7.6, using the Zetasizer Nano ZS device, Malvern Instruments, UK) of the coated beads from -25.8 ± 0.4 mV of the uncoated beads to -11.5 ± 0.3 mV of all the coated beads regardless the coating intensity (mean ± 95% CI for all coated beads).

### In-vivo capture efficiency measurements

The capture efficiency of six ascidian species was measured either underwater in their natural habitat (in situ) or under controlled conditions in a running seawater facility (laboratory experiments). These were *Microcosmus exasperatus* (laboratory experiments), *Herdmania momus* (laboratory experiments), *Styela plicata* (laboratory experiments), *Polycarpa mytiligera* (in-situ experiments), *Ciona intestinalis* (in-situ experiments) and *Styela clava* (laboratory experiments).

For in-situ experiments, inhaled and exhaled water were directly and simultaneously collected as described by (20). This non-intrusive approach, known as the VacuSIP-InEx method (44), was previously used to study the feeding and metabolism of suspension feeders in situ (19–21, 44). Briefly, two small sampling tubes were introduced into the openings of individual animals, and water was collected into evacuated vessels. We used the method of Jacobi et al. (20) to introduce artificial beads to the inhaled water. Briefly, polystyrene beads were delivered to the animals using a loop of a perforated tube placed around the inhalant siphon. Experiments were conducted at the Eastern Mediterranean (EMT), the Northern Red Sea, and the Northwest Atlantic (Long Island Sound; LIS).

To compare the efficiency by which ascidians capture the beads, paired samples of water at the inlet and outlet of the filtering system of ascidians were collected as described above. The capture efficiency was defined as:

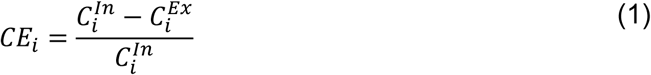

where *i* is the particle type and 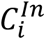 and 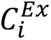 are the concentrations of type *i* particles in the inhaled and exhaled water (i.e., before and after filtration), respectively.

### Flow cytometry

Particle concentration in the inhaled and exhaled water samples was measured by flow cytometry as described by Jacobi et al. (20, 21). The instrument used (AttuneNXT® Acoustic Focusing flow cytometer, Applied Biosystems) allowed for high-precision measurement of particle concentrations (±5%). Samples were preserved with 0.1% Glutaraldehyde (final concentration) and kept at 4°C until they were analyzed (within two weeks). Samples were not frozen to prevent breaking and loss of the polystyrene beads.

### Diffusive and advective transport through the mucous filter

#### Mucus harvesting

The mean square displacement (MSD) of beads in fresh mucus was measured to quantify their mobility in the ascidian mucous filter. To obtain the mucus, the water supply to a beaker containing an individual ascidian was stopped, and a menthol solution (Sigma-Aldrich, cat. M2772 dissolved in absolute ethanol) was added to the beaker at a final concentration of 0.1 gr L^-1^. Once the animals stopped reacting to gently poking with a silicone swab (usually within 15 minutes), a swab was gently inserted into the inhalant siphon, and the mucus was gently harvested by gently swabbing along the inner walls of the branchial sack. After the harvest, the beaker was resupplied with seawater (17). In most cases, mucus harvesting did not harm the animals, and they resumed feeding at normal efficiency within an hour (17). Harvesting was done only after capture experiments were completed, and animals were not used for capture measurements again. Samples were pooled out from several ascidians to obtain enough mucus. No fixative agent was used, and mucus samples were kept chilled until MSD measurements were done (within 3 hours).

#### Diffusive transport - mean square displacement (MSD) measurement

Beads were embedded in mucus, and the mucus-bead mixture was transferred onto a microscope slide and mounted on the microscope (BX 73, Olympus LS, Japan). A 60-second video of the sample was recorded at 15 frames sec^-1^, then the slide was shifted to observe a new field of view, and another 60 seconds were recorded. Image analysis was made with the Fiji version of the ImageJ software (45) using the TrackMate plugin (46). The location of the beads in each frame (∼6 beads per video) was used to calculate the MSD of each particle across all the recorded time lags (τ). The data was used to construct an MSD versus time lag curve for each bead type. The slope of these curves was used to calculate the ‘effective diffusion coefficient’ (D_eff_) of each bead type. To obtain statistically robust data, the data obtained for τ > 30 was omitted. For additional details, see the supplementary material.

#### Advective transport – flow cell experiments

Particle transport through the mucous filter was done using a custom-built microfluidics setup containing a membrane filtration flow cell fitted with an optical window as described in (47). Fresh mucus was deposited on an ultra-filtration membrane as a structural support for the mucus. This membrane-mucus specimen was placed in the flow cell, and a pressure gradient was applied to it so that seawater (0.2 µm filtered) flowed through the system at ∼750 µL min^-1^. Then, at t = 0, two experimental bead types were added to the mucus at similar concentrations (particles mL^-1^). The particles used were fluorescent beads that contain different fluorophores. At each run, one type of bead was coated with a certain Pluronic (F68 or F108) while the other remained uncoated or coated with a different Pluronic. Four different regions were visualized with a confocal microscope (TCS SP8, Leica) every 5 minutes for a total of one hour to follow the advection of beads through the membrane-mucus system. Image analysis was done with an in-house MATLAB program. Each scan was converted to a gray-scale image. Then, based on their intensity, the pixels in each scan were categorized into ones that showed parts of a particle or empty pixels. The location of the supporting membrane was determined based on the laser reflection from the membrane surface. So, the surface coverage in % (i.e., the percentage of the observed surface covered by beads) as a function of the distance from the supporting membrane, z (Fig. 4E), was determined. Additional details, including a schematic illustration of the experimental setup, can be found in the supplementary material (Fig. S2).

### Characterization of the mucous filter

The carbon and nitrogen content of the mucous filter was measured using standard me CHN analysis with a PerkinElmer 2400 series II SHNS/O Analyzer (see Supplementary material for details).

Cryogenic scanning electron microscopy (cryo-SEM) was used to visualize fresh ascidian mucus. A suspension of uncoated beads was added to the water, and then animals were anesthetized with menthol, as described above, and dissected to expose the mucus that covered the branchial sac. The bead-laden mucus was visible on top of the branchial sac in all cases. Specimens of mucus-covered branchial sac tissue were frozen using liquid ethane and immediately imaged.

Proteomic analysis was done on five mucus samples, three water samples taken from the inner part of the branchial sac, and three branchial sac tissue samples. All samples were digested by trypsin and analyzed by liquid chromatography with tandem mass spectrometry (LC-MS/MS) using a Q-Exactive^TM^ mass spectrometer (ThermoFisher). Peptides were identified using the Proteome Discoverer software (version 1.4, ThermoFisher) using the Sequest search algorithm on *Ciona intestinalis* and *Ascidiacea uniport* databases. The peptide content and composition of the mucus, branchial sac tissue, and ambient water were tested for similarity. While mucus and tissue samples were found to be different from water samples collected from the branchial sac (ANOSIM, p<0.05), they did not significantly differ from each other.

### Statistical methods

Statistical analysis was done with the R language and environment for statistical computing (48) using the RStudio environment (Version 1.4.1106, © 2009-2021, RStudio). Differences in the capture efficiency of the beads were tested using a Repeated Measures Permutational ANOVA test (49). Post hoc analysis was carried out by a permutational pairwise t-test of logit transformed capture efficiency data with 1000 permutations for each pair of data, and *p* values were adjusted according to the Bonferroni method. Unless stated otherwise, data are presented as mean ± 95% confidence interval of the mean.

## Acknowledgments

We thank the staff of the Faculty of Civil and Environmental Engineering (Technion), Faculty of Marine Sciences in Michmoret (Ruppin Academic Center), the Inter-University Institute in Eilat, and the Department of Marine Sciences (UConn, U.S.A.) for support and use of the scientific diving centers and running seawater facilities. Special thanks to Aviv Ben-Tal, Keats Conley, Rei Diga, Noam Gridish, and all members of the Yahel, Ramon, and Shavit labs, Prof. Evan J. Ward (UConn, U.S.A), Dr. Na’ama Koifman and the staff of the Technion Center for Electron Microscopy of Soft Matter, and The Smoler Protein Research Center (Technion).

